# Human visual motion perception shows hallmarks of Bayesian structural inference

**DOI:** 10.1101/2020.11.05.370452

**Authors:** Sichao Yang, Johannes Bill, Jan Drugowitsch, Samuel J. Gershman

## Abstract

Motion relations in visual scenes carry an abundance of behaviorally relevant information, but little is known about the computations underlying the identification of visual motion structure by humans. We addressed this gap in two psychophysics experiments and found that participants identified hierarchically organized motion relations in close correspondence with Bayesian structural inference. We demonstrate that, for our tasks, a choice model based on the Bayesian ideal observer can accurately match many facets of human structural inference, including task performance, perceptual error patterns, single-trial responses, participant-specific differences, and subjective decision confidence, particularly when motion scenes are ambiguous. Our work can guide future neuroscience experiments to reveal the neural mechanisms underlying higher-level visual motion perception.

## Introduction

Motion relations in visual scenes are a rich source of information for our brains to make sense of our environment. We group coherently flying birds together to form a flock, predict the future trajectory of cars from the traffic flow, and infer our own velocity from a high-dimensional stream of retinal input by decomposing self-motion and object motion in the scene. We refer to these relations between velocities as *motion structure*.

In general, motion structure is not directly observable; it must be inferred from ambiguous sensory input. Bayesian inference provides a principled mathematical framework for dealing with this ambiguity. Understanding visual motion perception within a Bayesian framework has a long history in perceptual psychology, with applications to diverse phenomena: contrast-dependent speed perception (Ascher and Grzywacz, 2000), perceptual biases originating from partial occlusions of moving rigid objects (Weiss, Eero P Simoncelli, and Adelson, 2002), the perception of superimposed drifting gratings (Hedges, Alan A Stocker, and Eero P Simoncelli, 2011; Weiss, Eero P Simoncelli, and Adelson, 2002), depth perception of moving objects (Welchman, Lam, and Bülthoff, 2008), and speed perception of naturalistic stimuli (Chin and Burge, 2020). This previous work, however, was focused on elementary motion structures, in the sense that stimulus features either moved independently or were grouped to form rigid objects. Our goal is to understand how more complex and realistic forms of structure are inferred.

Many real-world motion patterns can be decomposed into hierarchical structures, where some motions are nested inside others (Gershman, Tenenbaum, and Jäkel, 2016). For example, the motion of flocking birds can be decomposed into shared and unique components. Intuitively, we see the motion of a bird in terms of its motion relative to the reference frame of the flock, which itself is perceived as a coherently moving object (below we will be more precise about what this means mathematically). Yet the concept of a *flock motion* is a latent variable that cannot be observed directly. In fact, coherent motion is one of the principal cues that the visual system uses to group visual features into objects (Spelke, 1990; Wertheimer, 1938). Thus, the computational puzzle is to understand how hierarchical structure can serve as both an organizing force, bringing order to the unruly circus of visual features, while also being inferred from those same features.

Building on the seminal work of Johansson, (1950), which first systematically studied hierarchical motion decomposition in displays of moving dots, subsequent studies formalized the idea (Clarke, Öğmen, and Herzog, 2016; Grossberg, Léveillé, and Versace, 2011; Shum and Wolford, 1983) and identified principles of how motion components are extracted when multiple decompositions are possible (DiVita and Rock, 1997; Gogel, 1974; Restle, 1979). Recently, Gershman, Tenenbaum, and Jäkel, (2016) proposed a Bayesian framework for explaining motion structure discovery, using probabilistic inference over hierarchical motion structures (they called *motion trees*). Subsequently, Bill et al., (2020) established experimentally that humans can exploit hierarchical structure when solving demanding visual tasks, such as object tracking and trajectory prediction. However, since participants were informed about the structure underlying the tasks, it remained unclear whether participants could infer this structure from sensory input.

Here, we address this question by means of two theory-driven psychophysics experiments: alternative forced choice tasks in which human participants reported the perceived latent structure underlying short dynamic visual scenes. Key to studying the specific contribution of motion structure to perception is precise control over all object velocities. To this end, we adopted the stimulus design from Bill et al., (2020), which generates trials stochastically while endowing the experimenter with full control over the joint velocity distribution of all visible objects. We were thereby able to eliminate several confounds that plague many traditional visual motion displays, such as informative spatial arrangement of objects or informative low-level stimulus statistics. As an additional benefit, the resulting visual scenes followed an analytically tractable generative model which, when inverted according to Bayes’ rule, afforded an ideal observer of Bayesian structural inference. The ideal observer enabled us to detect the behavioral signatures of structural inference that become apparent when the evidence is ambiguous.

In our first experiment, we show that a choice model, which wraps a small set of known perceptual imperfections around an otherwise ideal Bayesian core, predicted many facets of human responses with high fidelity. This included motion structure-specific task performance, distinct patterns of the confusability of certain structures, predictions of responses with single-trial resolution, individual differences between participants, and subjective confidence reporting. In our second experiment, we specifically targeted human perception of highly ambiguous scenes, which we generated by morphing continuously between two prototypical motion structures. The choice model again predicted human choice probabilities accurately. We conclude our study with a model comparison revealing the differential contributions of the model’s components to explaining human motion structural inference.

## Results

### Visual motion scenes with hierarchical structure

We adopted the hierarchical motion structure representation from Gershman, Tenenbaum, and Jäkel, (2016) and Bill et al., (2020), and the stimulus generation from Bill et al., (2020). Their main components with respect to the present work are summarized in the following.

#### Graph representation of hierarchical structure

In Experiment 1, we focused on covert motion relations between 3 colored dots, using the 4 motion structures illustrated in **Figure 1A**: independent (*I*), global (*G*), clustered (*C*), and hierarchical (*H*) motion. In the case of independent motion, there are no correlations between the dots’ velocities. The independence is characterized in the graph structure (gray background in **Figure 1A**) by the absence of shared ancestors between the 3 visible dots (colored nodes). The illustration next to the graph exemplifies the dot velocities in a typical scene. For global (G) motion, a single *latent motion source* (gray node in the tree) is added, which passes its velocity down to its descendants. In addition, each visible node features a smaller independent motion component. Global motion describes, for example, to first order the velocities for a flock of birds. In order to mathematically describe the coupling strength between the dots, we endow each source with a *motion strength λ* (vertical extent of graph edges), with larger values of *λ* describing higher average velocities (see Materials and Methods for the mathematical definition). In the illustrated global motion scene, the shared motion source is the dominant component to the dot velocities. The third motion structure is clustered (*C*) motion, in which 2 of the 3 dots share a latent motion source (purple node) while the 3rd dot moves independently. Finally, in hierarchical (H) motion, two clustered dots and the 3rd dot are nested under an additional, global moving reference frame. The fact that three is the smallest number of objects supporting such deeply-nested hierarchies is why we chose a 3-dot stimulus design.

**Figure 1.**
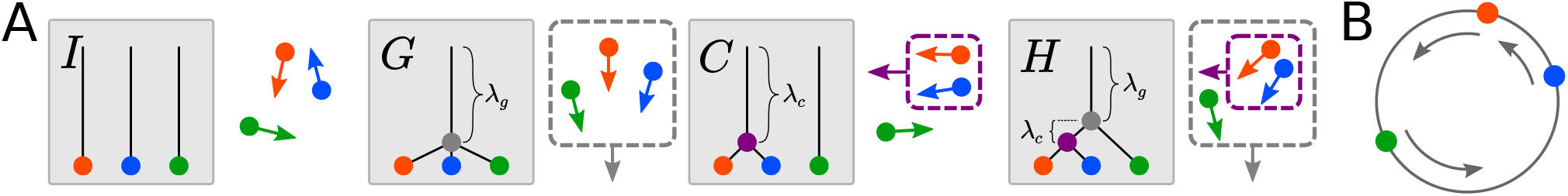
Motion structures used in Experiment 1. **(A)** Tree representations of 4 motion structures, each with 3 moving objects: independent (*I*), global (*G*), clustered (*C*), and hierarchical (*H*) motion. Next to each tree, an example of the resulting motion patterns is provided. The motion structures are unobservable (latent), but govern the velocity relations of the 3 visible objects (colored dots). In our formal representation of structure, object velocities are driven by latent motion sources (non-leaf nodes in the graph; dashed boxes in the examples) that affect the velocities of all connected downstream objects. Each motion source features a motion strength *λ* (vertical edge length in the graph) with larger strengths corresponding to higher average velocities. For example, the purple motion source drives the red and blue dots, either strongly (in structure *C*) or weakly (in *H*). **(B)** In the experiments, colored dots moved along a circular track in order to make spatial relations among objects uninformative. Dot colors were randomized in each trial. Here, a global (*G*) motion stimulus is illustrated.

#### Stochastic generation of visual scenes

The graphs in **Figure 1A** do not specify how a motion structure gives rise to a concrete dynamic visual scene. We created motion scenes for our experiments using the stochastic stimulus generation proposed in Bill et al., (2020); see Materials and Methods for details. In this stimulus design, dot velocities *v*(*t*) = (*v*_1_(*t*), *v*_2_(*t*), *v*_3_(*t*)) follow a multivariate Ornstein-Uhlenbeck process (Gardiner, 2009) in which latent motion sources play the role of random forces which accelerate or decelerate the dots. The resulting velocity trajectories feature the desired motion relations, obey our intuition that velocities change gradually due to inertia, and have an analytically tractable joint probability distribution. Analytical tractability enabled us to give all dots the same marginal speed distribution, independent of dot identities and the underlying motion structure. Thus, the velocity of each dot taken alone was entirely uninformative about the motion structure. In addition, all dots were placed on a circle, that is, *v*(*t*) describes their angular velocity (see **Figure 1B**). Circular stochastic stimuli have the advantage that they asymptotically break any spatial patterns between the dots, and thus leave correlations between velocities as the only information available for distinguishing different structures.

### Humans can identify latent motion structure, but feature distinct error patterns

Using the above-described stochastic motion scenes, we invited 12 participants to perform the visual motion structural inference task illustrated in **Figure 2A**. On each trial, a 4 s long dynamic scene with latent underlying motion structure *S* ∈ {*I, G, C, H*} was presented on a computer screen (see Materials and Methods for details, and **Supplemental Video 1** for a screen recording of several trials). The structure *S* was drawn uniformly before each trial, and remained fixed throughout the trial. After 4 s, the scene froze and the participant reported with a mouse click the perceived structure and their level of confidence (low/high) in the decision. After the decision was reported, the correct answer was revealed. **Figure 2A** exemplifies the feedback for a wrong choice (C), when the correct answer was (G). For a low confidence decision, the participant received +1 point for correct answers and 0 points for incorrect answers. For a high confidence decision, an additional point was “bet” which could be matched (correct answer) or lost (wrong answer). Participants were aware of the 4 structures, received brief training on the task, and conducted 200 trials (achieved score: 190 ± 33 points (mean ± SD)). In order to assess the reliability of responses, the 200 trials of each participant consisted of only 100 unique scenes, each presented twice in randomized order.

**Figure 2.**
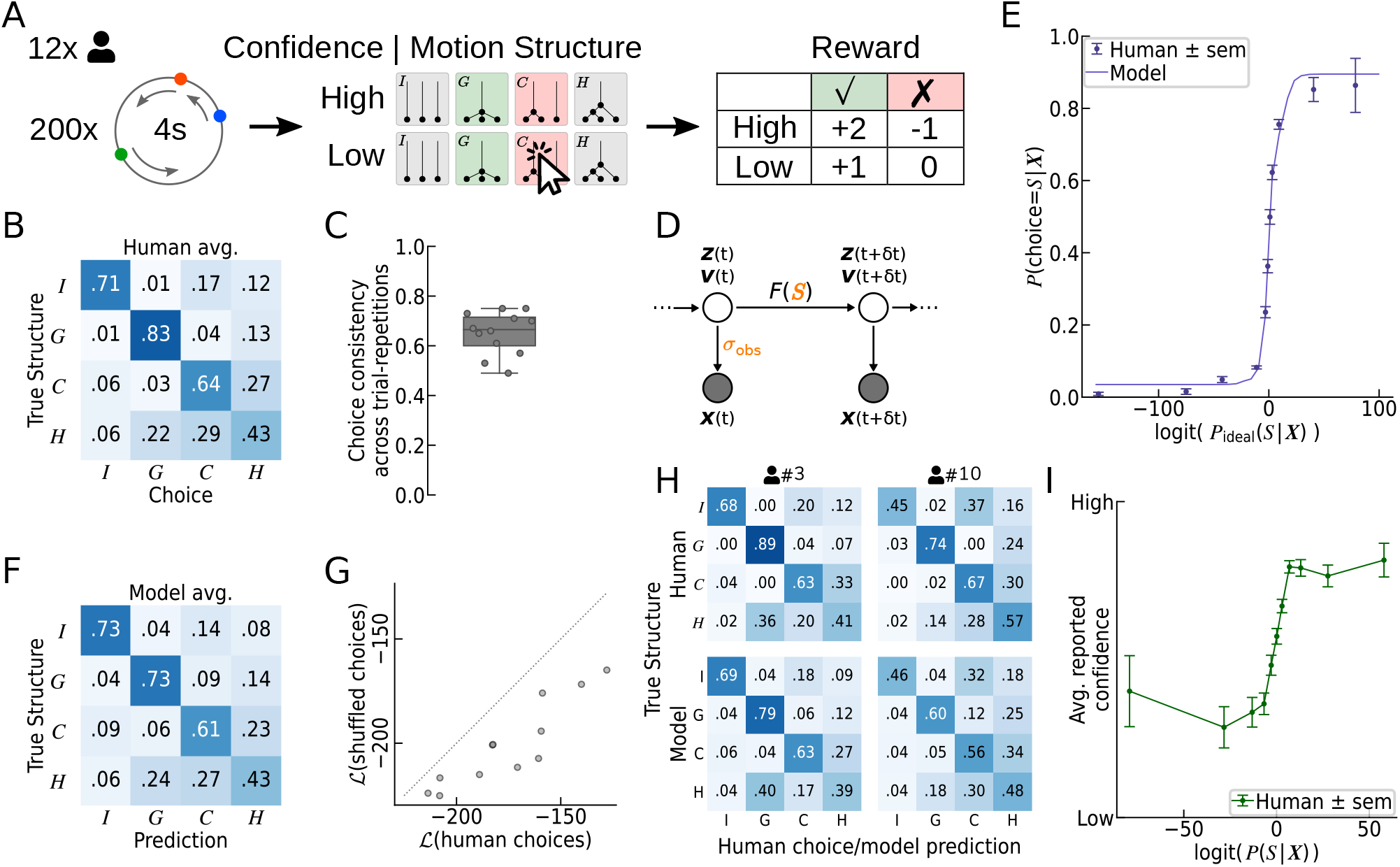
Human perception of structured motion shows hallmarks of Bayesian structural inference. **(A)** Illustration of Experiment 1. In each trial, a short scene of three moving dots was presented. The scene was generated from a latent structure drawn randomly from the 4 structures in **Figure 1A**. At the end of the trial, participants reported the perceived structure (*I/G/C/H*) and their confidence (low/high) in that decision. The correct structure was then revealed, and points were awarded based on correctness and confidence. **(B)** Confusion matrix of human responses, averaged over all participants and trials. The (*S,S′*) element is the empirical probability of choosing *S′* in trials of structure *S*. Humans perform clearly above chance (0.25), but show a reliable pattern of errors. **(C)** Each trial was presented twice to the same participant. Human choice consistency across repetitions was significantly below 1, revealing a stochastic component in human motion structural inference. Each dot represents one participant. Box plots mark the 0, 25, 50, 75, and 100 percentiles. **(D)** Generative model underlying the Bayesian ideal observer and the choice model. True dot locations, ***z***(*t*), and velocities, *v*(*t*), evolve stochastically with motion structure-dependent transitions ***F***(*S*). Only noisy observations of the locations, ***x***(*t*), are available to the observer. Kalman filtering inverts this generative model, while assuming different candidate motion structures S, and allows us to calculate the likelihood *p*(***X*** | *S*) that structure *S* generated the observed trajectory ***X***. **(E)** Psychometric function. Human responses show a sigmoidal transition as a function of the ideal observer’s log-odds. The log-odds measure the statistical evidence for each of the four structures *S* to have generated trial trajectory ***X***. Our fitted choice model of human responses predicts the psychometric function of the participants. Predictions were obtained via cross-validation to avoid over-fitting the data. **(F)** Confusion matrix of the choice model, averaged over all participants and trials (cf. panel B). The fitted model predicts human responses, including typical perceptual errors. **(G)** Predictions at single-trial resolution. A shuffled data set removed the stimulus–response correspondence while leaving the confusion matrix unchanged, for each participant. Refitting the model leads to significantly lower response likelihoods than for the non-shuffled human responses. Each dot represents one participant. Two participants’ points are overlapping. **(H)** Confusion matrices of two representative participants and their respective model predictions. Participants featured different error patterns, which the model recapitulates. **(I)** Subjective confidence reports show signatures of Bayesian inference. The average reported confidence is a sigmoidal function of the posterior log-odds, logit(*P*(*S* | ***X***)).

Our participants had a classification accuracy well above chance level for all four motion structures, as evidenced by the confusion matrix shown in **Figure 2B**. Yet, the different motion structures elicited distinct error patterns. For example, participants tended to confuse clustered (*C*) and hierarchical (*H*) stimuli, or perceived hierarchical motion as global (G). Interestingly, the confusability of motion structures often was not symmetrical, as exemplified by the *I-C* and *C-I* elements of the confusion matrix. The spatial arrangement of the dots on the circle, in contrast, did not seem to have an influence on the perceived motion structure (see **Supplemental Figure S1**), in line with the uninformativeness of spatial arrangement in the task design. Furthermore, human responses featured a truly stochastic component, as revealed by comparing their responses across the two unique-trial repetitions (**Figure 2C**). When re-presented with an identical scene, the same participant’s choice was consistent with their previous choice only 65.2% ± 1.4% (s.e.m.) of the time. To summarize, our participants were generally able to identify the structure underlying a motion scene. Yet, their responses featured stochasticity and systematic error patterns, with some structures being more confusable than others.

### Human motion perception is explained by a model of Bayesian structural inference

Erroneous identification of motion structures could arise either from false perceptual decision making or— owing to the stochastic stimulus generation—from objective ambiguity of the presented scenes. To tease these factors apart, we analyzed human responses from a normative standpoint of statistical inference. Concretely, we progressed in two steps. First, we exploited the analytical tractability of the stimulus design to formulate a *Bayesian ideal observer*. The ideal observer evaluates the likelihood *p*(***X*** | *S*) of a presented trajectory, ***X***:= {***x***(*t*), 0 ≤ *t* ≤ *T*}, under the four alternative motion structures *S* and, thus, provides an objective statistical measure of a trial’s most likely structure and of the statistical significance of this conjecture. In the second step, we devised a *choice model* of human choices, *P*(choice = *S* | ***X***), by augmenting the ideal observer with well-known imperfections of human perceptual decision making (see Materials and Methods). The choice model featured a small number of interpretable free parameters and afforded a deeper understanding of the computations driving human motion structural inference. Both models and their implications are presented in the following.

#### The Bayesian ideal observer suggests statistically correct structural inference by humans

The analysis via an ideal observer was made possible by the analytical tractability of the stimulus that obeyed the generative model shown in **Figure 2D**. True dot locations ***z***(*t*) and velocities ***v***(*t*) are latent variables that evolve according to a stochastic process with correlated transitions ***F***(*S*), which depend on the underlying latent structure *S*. For instance, a global motion stimulus features positively correlated transitions of dot velocities. Ultimately, the correlated transitions lead to temporal correlations among the observed locations ***x***(*t*), which we assumed to be noisy observations of ***z***(*t*) at the times of the video frames, *t* = 0, *δt*, 2*δt*,.., *T*. The linear stochastic dynamics of the generative process render Kalman filtering with a correlated state-transition model the ideal observer for these stimuli (see Materials and Methods). In particular, Kalman filtering allows us to calculate the likelihood, *p*(**X** | S), of a trial’s trajectory under any of the candidate motion structures, *S* = *I, G, C, H*. Thus, comparing the likelihoods *p*(***X*** | *S*) across candidate structures provides an objective statistical measure not only of a trial’s most likely motion structure, but also of its ambiguity.

Using the ideal observer, we evaluated human choices on all trials in terms of the log-odds, log*P*_ideal_ (*S* | ***X***) – log*P*_ideal_ (not *S* | ***X***), where *P*_ideal_ (*S* | ***X***) « *P*(*S*) *p*(***X*** | *S*) is the ideal observer’s posterior belief over the four structures *S*, using a uniform prior 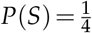. The frequency of human choices, *P*(choice = *S* | ***X***), as a function of the ideal observer’s log-odds is shown in **Figure 2E**. The psychometric function exhibited the sigmoidal shape expected from Bayesian decision making, and, thus, pointed to an element of correct statistical reasoning by our participants over the latent structural features underlying the presented scene.

#### A model of human choices: wrapping imperfections around a Bayesian core

Inspired by the psychometric function (**Figure 2E**) and the stochasticity of human responses (**Figure 2C**), we devised a choice model that added well-known imperfections of human perceptual decision making to the Kalman filter’s Bayesian core (Acerbi, Vijayakumar, and Wolpert, 2014; Bill et al., 2020; Drugowitsch, Wyart, et al., 2016; Wichmann and Hill, 2001):

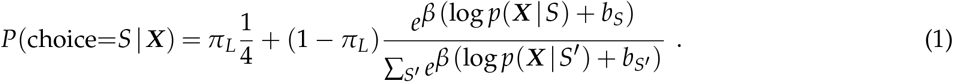

The model has a small lapse probability, *π_L_*, for guessing randomly, three a priori-biases, *b_S_* with *S* ∈ {*G, C, H*}, playing the role of a log-prior 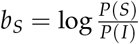 relative to the independent structure, and an inverse temperature, 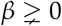, for adjusting the amount of decision noise. One additional parameter is the observation noise, *σ*_obs_, that enters the likelihood *p*(***X*** | *S*) through the Kalman filter. For better identifiability, we fitted the lapse probability *π_L_* and the observation noise *σ*_obs_ globally to all participants, leaving four free parameters (*β, b_G_, b_C_, b_H_*), which we fitted per-participant via maximum likelihood (see Materials and Methods). Thus, in the choice model, the only connection between the presented trial ***X*** and the predicted choice ***S*** is formed by the Bayesian core log *p*(***X*** | *S*) from the Kalman filter.

#### Prediction of human choice patterns

The choice model in Equation 1 predicted human responses across all motion conditions, as evident from comparing the model’s confusion matrix in **Figure 2F** with the confusion matrix of human participants (**Figure 2B**). Besides the qualitative similarity of responses, the model also captured the “fine-structure” of typical human errors, such as the asymmetry between the I-to-C and C-to-I elements. Notably, the model’s confusion matrix was obtained via leave-one-out cross-validation. This makes the match to human responses a prediction, not a result of over-fitting. Furthermore, the model replicated the psychometric curve in **Figure 2E** with high accuracy, and featured response consistencies across trial-repetitions that were qualitatively similar to human responses (see **Supplemental Figure S2**).

#### Prediction of human choices at single-trial resolution

The confusion matrix does not contain information on the exact trial that a certain response had been given to. We wondered if the model’s predictions only matched “summary statistics” of human responses, as expressed by the confusion matrix, or whether the model also captured responses to individual trials. To test this, we created a shuffled data set for each participant by randomly swapping the human responses to trials of the same motion structure. Thus, the shuffled data removed the stimulus–response correspondence under the constraint of leaving the confusion matrix unchanged. We then re-fitted the choice model to the shuffled data. As shown in **Figure 2G**, the model explained the original, non-shuffled data better than the shuffled data, for all 12 participants (cross-validated). This suggests that the model captured features of human motion structural inference at trial-level resolution.

#### Prediction of individual differences between participants

When analyzing the data of different participants, we noticed participant-specific differences in their confusion matrices (see top row of **Figure 2H** for two representative participants). Differences were particularly prominent for perceiving independent motion as being clustered (I-C element) or whether hierarchical motion was preferentially confused with global or clustered motion (H-G and H-C elements). The model can account for such perceptual biases via the prior parameters *bS*. We found that the biases allow the model to feature the same participant-specific differences as the participants, as can be seen by the predicted confusion matrices (bottom row of **Figure 2H**; see **Supplemental Figure S3** for the data of all participants). In principle, the individual differences observed in **Figure 2H** could have arisen from the stochastic trial generation since each participant had been presented with a different set of stimuli. To rule out such stimulus-induced effects, we compared how well each participant’s model predicted the responses of all other participants, confirming the unique fitness of each participant’s model parameters (see **Supplemental Figure S4**).

Overall, the choice model’s ability to predict various aspects of human responses supported the hypothesis that human motion structural inference could be understood as probabilistic inference over the presence (or absence) of latent motion features.

#### Human subjective confidence shows signatures of Bayesian decision confidence

Beyond the reported motion structure, we asked if our participants’ subjective confidence was dependent on the stimulus ambiguity, as measured by the Bayesian model. The relation between experimentally reported subjective confidence and theoretically derived Bayesian confidence has been studied for a variety of tasks (Drugowitsch, Moreno-Bote, and Pouget, 2014; Galvin et al., 2003; Hangya, Sanders, and Kepecs, 2016; Kepecs and Mainen, 2012; H.-H. Li and Ma, 2020; Mamassian, 2016; Pouget, Drugowitsch, and Kepecs, 2016; Sanders, Hangya, and Kepecs, 2016), and the literature is often equivocal about the exact nature of the relation. We analyzed the participants’ reported confidence as a function of the Bayesian predicted confidence, *p*(*S* | ***X***), where the prior *P*(*S*) is given by the fitted biases *b_S_* for each participant, and the structure *S* is evaluated at the human choice (Pouget, Drugowitsch, and Kepecs, 2016). Thus, the Bayesian predicted confidence expresses the subjective belief in having made the correct decision, under the assumption that participants calculate their subjective posterior over the four structures (without decision noise or lapses). In **Figure 2I**, the participants’ reported confidences are shown as a function of the log-odds of the Bayesian predicted confidence, logit(*P*(*S* | ***X***)). The logistic shape of the psychometric curve indicates that participants employed an estimate of their subjective perceptual (un-)certainty, and judged their own decision in line with Bayesian models of confidence. Also, when evaluating the confidence in terms of the ideal observer’s posterior—that is, disregarding putative subjective priors *p*(*S*)—the psychometric function showed an approximately monotonic increase (see **Supplemental Figure S5**). Thus, our data suggest that the mental computation of confidence obeys principles of correct statistical reasoning.

### Hallmarks of Bayesian structural inference when perceiving ambiguous scenes

Signatures of statistical reasoning become particularly apparent in the face of ambiguous stimuli. To put our hypothesis—that human motion structural inference can be understood as Bayesian inference—to test, we focused on a pair of motion structures which had been identified in pilot data to be particularly confusable: clustered (C) and hierarchical (H) motion. In Experiment 2, which directly followed Experiment 1, the same participants reported their percept in a two-alternative forced choice (2-AFC) task which specifically asked them to distinguish between clustered and hierarchical motion (see **Figure 3A**). The task was designed similar to Experiment 1, with two notable differences. First, participants did not receive any feedback about the correctness of their choices, in order to prevent further learning. Second, unbeknown to the participants, we presented not only stimuli of the C- and H-structure, but also stimuli of intermediate structure, i.e., stimuli of particularly high ambiguity (see x-axis of **Figure 3B**). Such morphing between motion structures was made possible by our tractable structure representation, which allowed us to gradually introduce a global motion component of strength *λg* while simultaneously reducing the strength *λc* of the clustered component. A small value of *λg*, for example, entails a barely discernible global correlation in velocities. Importantly, the morphed structures did not alter the marginal speed distributions of the dots, thus leaving dot velocity correlations as the only feature to build the percept on (see Materials and Methods for details on the experiment design). Experiment 2 was therefore tailored to investigating motion structural inference of highly ambiguous scenes.

**Figure 3.**
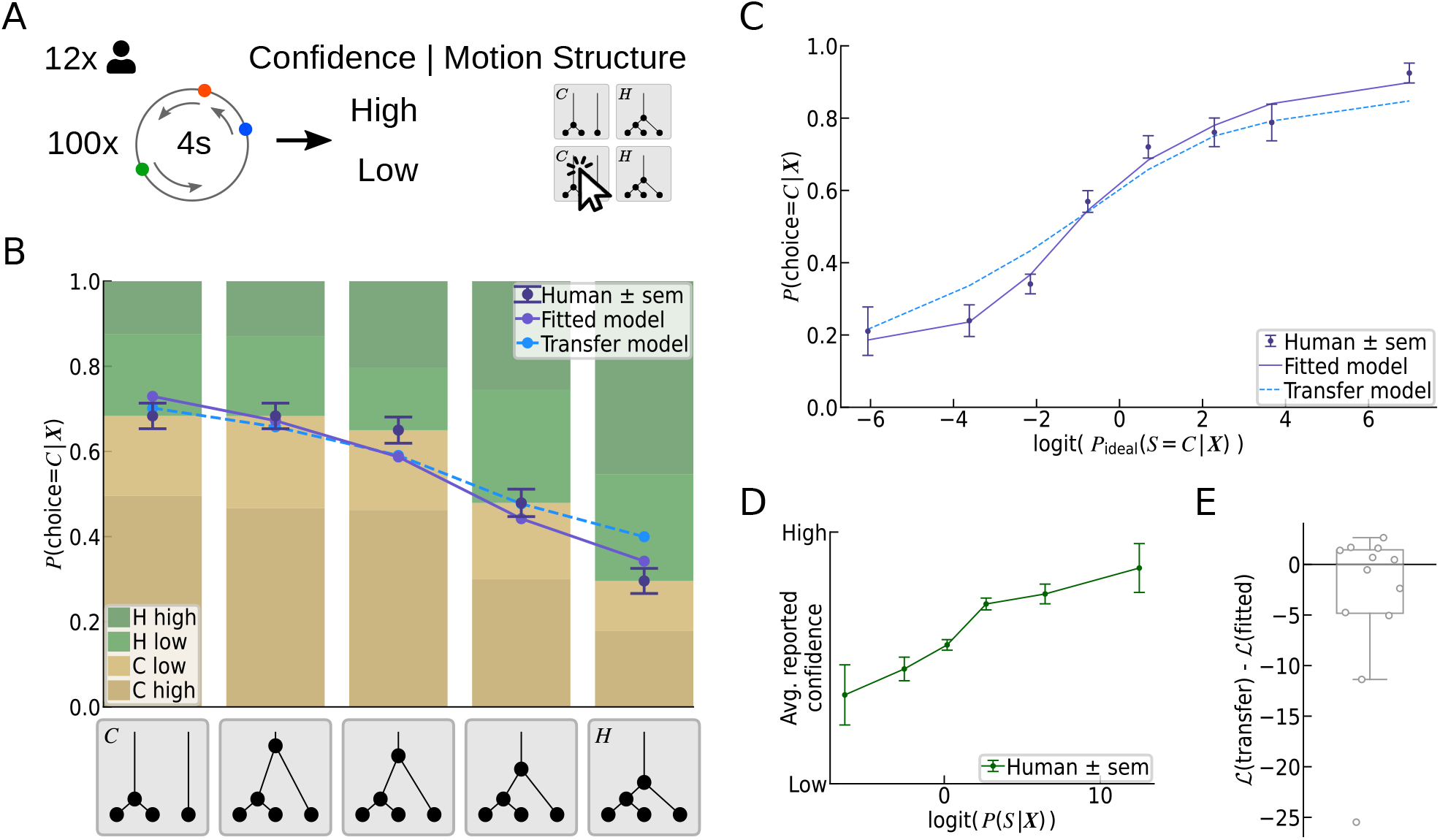
Predicting human motion structural inference of ambiguous scenes. **(A)** Illustration of Experiment 2. In a 2-AFC task without feedback, participants had to distinguish between the especially confusable *C*- and *H*-motion structures. **(B)** Gradual transition of choice probabilities. *Bottom:* unbeknown to the participants, we presented also stimuli of intermediate structures by morphing between the C- and H-prototypes. The resulting ambiguous scenes specifically probed for signatures of statistical evidence integration in human motion perception. *Top:* Human choices gradually shifted with changing structure (purple data points). The choice model, with fitted temperature and bias for each participant, predicted human choice probabilities accurately (purple line; cross-validated). Also the model from Experiment 1, termed the “transfer model”, qualitatively predicted human choices without refitting any parameters (dashed blue line). Bar plots: human choices (C: ocher, H: green) and decision confidence (color intensity). **(C)** Psychometric function. Human choices, the fitted choice model, and the transfer model, all exhibit a sigmoidal transition as a function of the ideal observer’s log odds. The fitted model’s predictions were obtained via cross-validation to prevent over-fitting. **(D)** Human reported confidence follows a monotonically increasing (sigmoidal) curve as a function of the log-odds of the Bayesian predicted confidence. **(E)** The cross-validated log-likelihood difference between the transfer model and the fitted model suggests that most participants are explained equally well by both models.

As expected for a Bayesian observer, and as shown in **Figure 3B**, human reporting gradually shifted from clustered (C) to hierarchical (H) percepts as the underlying structure moved between the two prototypes. Furthermore, human responses followed the signature logistic shape when trials were evaluated in terms of an ideal observer’s log-odds (see **Figure 3C**). Analogous to Experiment 1, we subsequently fitted our choice model to each participant. For the 2-AFC task, the model had only two free parameters, namely, inverse temperature *β* and bias *b_H_* for each participant (we maintained the shared parameters *π_L_* and *σ*_obs_ from Experiment 1). The model, shown as purple lines in **Figure 3B & C**, predicted human responses accurately, thereby corroborating our hypothesis of structural inference in the face of uncertainty.

#### Predictability of human confidence reporting

As for Experiment 1, we asked how the subjective confidence of our participants varied with the (un-)certainty of the presented motion scenes. As shown in **Figure 3D**, human confidence reporting followed a sigmoidal curve as a function of logit(*P*(*S* | ***X***)), as expected by models of Bayesian decision confidence (Pouget, Drugowitsch, and Kepecs, 2016).

### Generalization of model predictions across both experiments

While human responses in both experiments were accurately described by the separately fitted choice models, we wondered if model predictions would generalize across both experiments. To test for generalization, we evaluated, for each participant, the trials of Experiment 2 using the model fitted to Experiment 1 (excluding the unavailable ‘I’ and ‘G’ responses). We refer to this model as the *transfer model*. Predicting human choices across the two experiments involves two types of generalization: reduction to a smaller choice set, and the ability to make sense of “morphed” motion structures, which had neither been part of the transfer model’s nor of the humans’ training set (i.e., Experiment 1).

The predictions of the transfer model are shown as dashed, light-blue lines in **Figure 3B & C**. While not reaching the accuracy of dedicated model fits, the transfer model’s predictions of human choices were qualitatively correct. Concretely, the transfer model featured a shallower slope, but did not show any significant bias. Thus, the model appeared to generalize, at least qualitatively, across the two experiments.

To corroborate this observation, we strove to compare both models at the level of individual participants. Such comparison was complicated by the small number of trials in Experiment 2. We had limited the number of trials to 100 per person for not risking the participants’ task engagement (there was no feedback), which led to only 20 trials per tested intermediate motion structure and participant. Accordingly, when evaluated for each participant separately, both human responses and the models’ predictions showed strong variability (see **Supplemental Figure S6**). We therefore employed cross-validated log-likelihood comparison in order to test whether the improved prediction accuracy of the directly fitted model warranted the additional free parameters involved in the fitting process. The log-likelihood of held-out trials is shown in **Figure 3E** for the transfer model relative to the fitted model. Half of the participants were actually slightly better described by the transfer model than by a newly fitted model, an effect likely attributed to the limited number of trials available for fitting the model in Experiment 2. Thus, while heterogeneous across participants, the cross-validated log-likelihoods indicate that the refitting of parameters does not warrant the added model complexity, in our experiment. In summary, our findings point to some generalization of the choice model across experiments, yet a study with more statistical power would be required to solidify this observation.

### Importance of different model components to explaining human responses

We conclude our study of visual motion structural inference with a systematic model comparison to assess which model components are essential for explaining human responses. To approach this question, we re-analyzed the data from the 4-AFC experiment of **Figure 2** by means of the data log-likelihood and the protected exceedance probability. In particular, we studied the importance of the choice model’s “Bayesian core” given by the Kalman filter.

#### Differential comparison of model components

We measured each model component’s importance by differentially removing it from the choice model and refitting the resulting reduced model. The log-likelihood ratio relative to the full model is shown in **Figure 4A** for models without lapses (*π_L_* = 0) or biases (*b_S_* = 0). We did not include a model with zero temperature, *β* → ∞, since the intrinsic stochasticity of human choices (cf. **Figure 2C**) rules out a deterministic model. We found that, for most participants, the full model explained human responses significantly better than the reduced models. For the model without lapses, the analysis highlights the importance of including a lapse probability (even if it is a shared parameter, as in our case). For the model without biases, in contrast, the reduced log-likelihood is no surprise since this model is contained within the full choice model as a special case.

**Figure 4.**
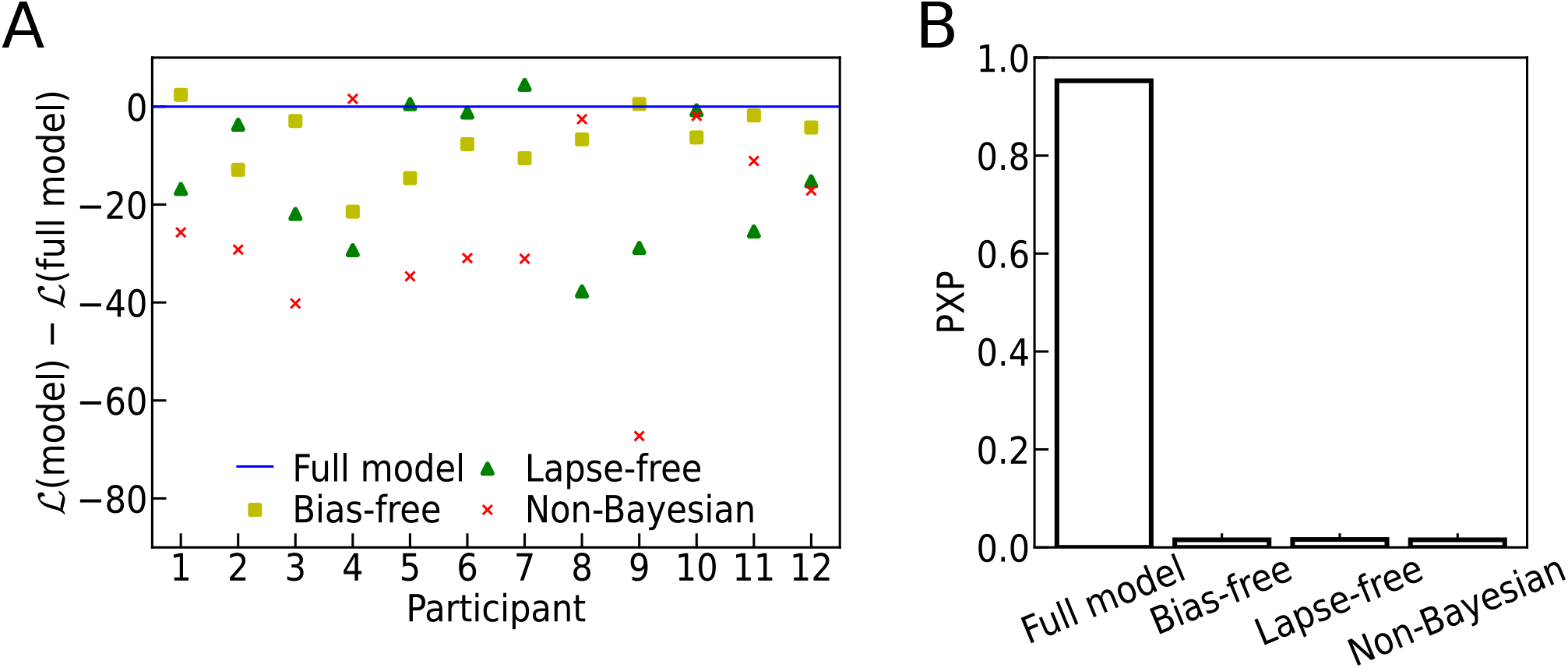
Importance of different model components to explaining human responses (Experiment 1). **(A)** Model comparison in terms of the cross-validated log-likelihood of human responses under different choice models with differentially replaced model components. The alternative models were re-fitted. All log-likelihoods are evaluated relative to the full model of Equation 1. For most participants, the full model explains human responses better than models with a uniform prior (all *b_S_* = 0, “bias-free”), without lapses (*π*_L_ = 0, “lapse-free”), and a model without the Bayesian core, *p*(***X*** | *S*), of the Kalman filter (“non-Bayesian”). **(B)** Bayesian model comparison in terms of the protected exceedance probability (PXP), using the cross-validated log-likelihood approximation to the model evidence. Also the PXP, which estimates the probability of a model to be the “winning” model, identifies the full model as prevalent over the alternatives.

Beyond relative model comparison, the log-likelihood of human choices provides an absolute measure of how well each model explains human responses (see **Supplemental Figure S7** for an evaluation relative to random choices). We find that some participants are generally better explainable than others. Indeed, it turns out that the full model’s log-likelihoods are highly correlated with the participants’ consistency across the two trial repetitions (cf. **Figure 2C**): more consistent participants’ responses have a higher likelihood under the model (Spearman’s rank correlation *ρ =* 0.91). This correlation is reassuring since it confirms that the prediction quality of the model strongly co-varies with the general predictability of the participants.

#### A non-Bayesian choice model explains human responses significantly worse

While the choice model built around the Bayesian core was sufficient for predicting many facets of human motion structural inference, we wondered if a Bayesian core was also necessary. In order to obtain at least a tentative answer to this question, we replaced the model’s Bayesian core—the stimulus log-likelihood log*p*(***X*** | *S*) obtained from Kalman filtering—by a non-Bayesian, yet reasonable, statistic: the Pearson cross-correlation matrix between all dot velocities. Concretely, we calculated for each trial ***X*** the correlation matrix Corr(***V***(***X***)) for the empirical velocities obtained from subsequent stimulus frames, 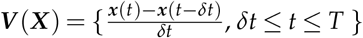. To make this measure as accurate as possible, we removed any putative observation noise by setting *σ*_obs_ = 0. The trial’s correlation matrix Corr(***V***(***X***)) was then compared to the expected correlation Corr(***V***(*S*)) under the different motion structures *S*, which can be calculated analytically for our stimuli (see Materials and Methods and **Supplemental Figure S8**). The negative distance to the prototypical correlation matrices, –║Corr(***V***(***X***)) – Corr(***V***(*S*))║_*r*_, measured with regard to an *ℓ_r_*-norm, replaced the Bayesian core in the model of Equation 1. We then re-fitted the four free parameters (*β, b_G_, b_C_, b_H_*) for each participant and the shared lapse probability *π_L_* and norm parameter *r*, for a fair comparison. The resulting non-Bayesian choice model, which resembles the generalized context model (Nosofsky, 1986), captured some aspects of human motion structural inference (see **Supplemental Figure S7**), a reassuring confirmation that the model was based on a reasonable statistic. It did, however, achieve a significantly lower quality-of-fit than the model with a Bayesian core, as evidenced by the model log-likelihood ratios in **Figure 4A**.

As a final check, we calculated the protected exceedance probabilties (PXP) for all considered models (Rigoux et al., 2014; Klaas Enno Stephan et al., 2009). The PXP estimates the probability of a model to be the most frequent model in a group of participants. The PXP is shown in **Figure 4B** and corroborates all earlier analyses by clearly favoring the full model over all tested alternatives. In conclusion, all considered components appear to contribute significantly to our choice model’s explanatory power—especially its Bayesian core.

## Discussion

We have shown that human observers can identify hierarchically nested, latent motion relations in visual scenes, and that they do so in close correspondence with Bayesian structural inference. Our choice model with a Bayesian core predicted a wide range of human responses across two psychophysics experiments. Besides general performance levels and typical error patterns (confusion matrices), the model captured participant-specific differences, trial-specific responses, subjective confidence, and the perception of the highly ambiguous motion scenes in Experiment 2 which neither humans nor the model had been trained on before. Key to isolating the contribution of motion structure to visual perception was an analytically tractable task design that rendered the presented scenes indistinguishable with regard to low-level stimulus statistics, such as spatial arrangement and marginal velocity distributions. Task tractability further afforded the formulation of an ideal observer via Kalman filtering, the Bayesian core for predicting human percepts. Notably, replacing this ideal core by a non-Bayesian statistic of velocity correlations led to a significantly reduced prediction quality. Together with the results of Bill et al., (2020), our findings suggest that the human mind employs a dynamical model of object motion, which enables it to identify and exploit latent structure in demanding visual tasks.

To investigate the isolated contribution of motion structure to visual perception, several simplifications had to be made in the stimulus design and analysis. In real-world scenes, motion relations are only one feature contributing to perception which is combined with other features such as the spatial arrangement, shape, texture and color of objects. Furthermore, the combinatorial complexity of possible structures underlying real-world scenes can hinder uncovering the computations driving structural inference. To make structural inference accessible to a quantitative treatment in the laboratory, we employed an impoverished stimulus that capitalized on motion relations as the only source of information and reduced the structural complexity to four categories. Nonetheless, the stochastically generated scenes of our study extend the complexity of traditional visual displays, such as drifting gratings or plaids, by supporting hierarchically nested motion structure and a wide range of dot trajectories.

The observed stochasticity of human responses to identical scenes (trial repetitions) is in line with previous studies (Acerbi, Vijayakumar, and Wolpert, 2014; Bill et al., 2020; Drugowitsch, Wyart, et al., 2016; Wyart and Koechlin, 2016), which we were able to partially capture with generalized probability matching, by raising the posterior to the power of *β*. The fact that our choice model shows a systematically higher stochasticity than human observers (cf. **Supplemental Figure S2**), however, is most likely attributed to additional mechanisms in human perception that are not captured by the model. One possible class of missing mechanisms are sequential effects (Cross, 1973; Petzschner, Glasauer, and Klaas E Stephan, 2015), that is, the stimulus of and response to one trial alters our percept of subsequent trials. In contrast, we assumed that each trial is processed in isolation for better tractability of the analysis.

Another experimental limitation with regard to a mechanistic understanding of structural inference is that our task design cannot separate multiplicative errors during sensory integration in the log-likelihood term of Equation 1 from stochasticity in the decision process. Both mechanisms are intertwined in the inverse temperature, *β*, in our model. An in-depth discussion of how to distinguish between these mechanisms can be found in Drugowitsch, Wyart, et al., (2016).

Finally, our model systematically under-predicted the identifiability of the global motion condition (cf. **Figure 2B+F**). Interestingly, a similar discrepancy was observed in Bill et al., (2020), where human performance exceeded the model’s predictions specifically in this motion condition. Globally coherent motion arguably is the most prevalent motion structure encountered by our visual system, triggered for instance by every saccade of our eyes. Evolution may have found solutions to process global motion more accurately than others structures.

Bayesian models of statistically ideal motion integration have been employed successfully to explain human motion perception for reduced stimuli in the laboratory (Ascher and Grzywacz, 2000; Alan A. Stocker and Eero P. Simoncelli, 2006; Weiss, Eero P Simoncelli, and Adelson, 2002) (but see Hammett et al., (2007)). However, for the complexity of real-world scenes, exact inference in any generative model, which would be sufficiently expressive to capture fine details, becomes intractable. Hierarchically structured models provide a strategy for the brain to tackle intractability by “abstracting away” behaviorally irrelevant details while facilitating inference over behaviorally relevant variables (e.g., the flock’s velocity in the example of observing many birds). One would expect that the expertise of which structural features are behaviorally most relevant evolves, at least in part, postnatally from experience. Indeed, developmental studies have found that structured motion perception begins to develop during the first year of infancy (Agyei et al., 2015) and continues to mature throughout later childhood (Ellemberg et al., 2004). Together with other studies, this led to the conclusion that *“motion processing systems detecting locally uniform motion mature earlier than do systems specializing in complex, globally non-uniform patterns of motion”* (Gilmore et al., 2007). Our work extends this line of research by demonstrating that adult participants can recognize nested, rotating motion relations.

The behavioral results of our study can furthermore inform new directions in neuroscience research. Cells in the middle temporal (MT) area show tuning to the local speed and direction of visual stimuli (Born and Bradley, 2005) and undergo temporal switching between representations of locally coherent velocity when input is ambiguous (K. Li et al., 2016). Neuronal firing in downstream areas, such as the dorsal medial superior temporal (MST) area, has been shown to carry information about elementary motion features, such as expansion or rotation (Graziano, Andersen, and Snowden, 1994; Mineault et al., 2012). To measure the brain’s representation of richer motion structures, it will be necessary to dissociate stimulus-induced neural responses from internally generated activity patterns that could reflect beliefs about structure. If adapted for animal experiments, our study could help link the values of task-relevant internal variables to neural recordings, particularly when the stimulus structure is ambiguous. With regard to computational modeling, we propose to investigate how on-line algorithms could implement the required Bayesian computations and how they could tame the combinatorial complexity of real-world motion structures.

## Materials and Methods

### Matrix representation of hierarchical structures

All motion structures in this study can be described as 3 objects driven by 5 motion sources: the global component (g), the cluster component (c), and three individual components (1,2,3), if we allow the global/cluster component to have zero strength in some of the structures. For example, the global motion source has zero strength in the clustered (C) condition. Bill et al., (2020) proposed that the hierarchical motion structures *S* from **Figure 1A** can be mathematically formulated as a matrix, combining a basis set ***B*** of motion features and the strength *λ* of each feature:

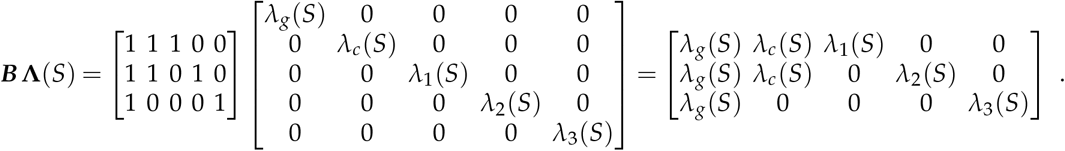

In 3 × 5 matrix ***B***, *B_k,s_* = 1 indicates that object *k* is driven by motion source *s*, and *B_k,s_* = 0 if object *k* is not affected by *s*. The basis ***B*** is shared across all used motion structures. The 5 × 5 diagonal matrix **Λ**(*S*) describes the structure-specific component strengths. The values of **Λ**(*S*) used in the experiments are provided in the experiment details, below.

### Stochastic generation of visual scenes

Dot velocities were generated stochastically from a multivariate Ornstein-Uhlenbeck process. Concretely, the dynamics of dot locations ***z***(*t*) and velocities ***v***(*t*) are given by

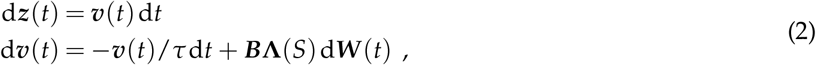

with 5-dimensional Wiener process ***W***(*t*) and friction time constant *τ* = 1.5s, the typical time scale of significant changes in velocity. In the discrete-time computer implementation, we updated the dot velocities and locations via the Euler-Maruyama method with time interval d*t* = 1ms. To stay on a circle, we constrained ***z***(*t*) to (*z*(*t*) mod 2*π*) after each integration step.

A stochastic dynamical system following Equation 2 converges to the stationary distribution

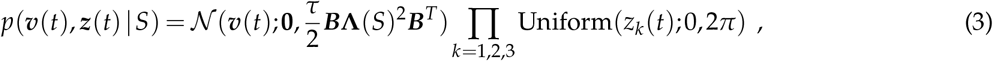

under the condition that all dots have a nonvanishing individual motion component, i.e., 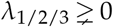 (see Gardiner, (2009)). This means that, in equilibrium, dot velocities are distributed according to a multivariate normal distribution and dot locations become asymptotically independent. When generating trials, we sampled the initial dot states from this stationary distribution in order to always stay in equilibrium. By choosing **Λ**(*S*) appropriately, we can keep the velocity distribution of each dot, i.e. each diagonal element of 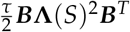, identical across all tested motion structures *S*, leaving velocity correlation as the only evidence for structural inference. We achieved this by ensuring *λ_g_*(*S*)^2^ + *λ_c_*(*S*)^2^ + *λ*_1_(*S*)^2^ = *λ_g_*(*S*)^2^ + *λ_c_*(*S*)^2^ + *λ_2_*(*S*)^2^ = *λ_g_*(*S*)^2^ + *λ*_3_(*S*)^2^ = 2^2^ throughout the study.

### Experiment details

12 Greater Boston residents (9 female, age 27.8±5.0) with normal/corrected-to-normal vision participated in the experiments in anechoic experiment rooms at Harvard for a $10 base pay and a $5.2 ± 0.9 performance-dependent bonus. After receiving brief verbal introduction to the experiment procedures and the four motion structures, participants sat in front of a 14” ThinkPad T460p laptop (viewable area: 31 cm x 18 cm, resolution: 1920 x 1080, refresh rate: 50 fps, operating system: Ubuntu 18.04). While viewing distance, viewing angle, head- and eye movement were unconstrained, participants viewed the screen perpendicularly from approximately 40 cm distance. Each participant completed three parts of motion structure identification trials: training, 4-AFC (Experiment 1), and ambiguous 2-AFC (Experiment 2). Trials across all parts shared the same computer interface. At the beginning of each trial, 3 dots in red, green, and blue (diameter: 0.6°of visual angle) were initialized on a black circle (diameter: 12° of visual angle) indicating the orbit of motion trajectories. To avoid color-based biases and heuristics, we randomly permuted dot colors for each trial. Initially, the dots were frozen for 0.5s, then moved for 4s following Equation 2, and then froze again for decision making. A 2×4 grid of buttons appeared at the lower right corner on the screen, and the mouse cursor appeared at the center of the screen, waiting for the participant to click a button according to their perceived structure (*I/G/C/H*) and decision confidence (high/low). Finally, decision feedback, trial count, and a brief instructional text were shown, guiding the participant to initiate the next trial with a mouse click or key press. Participants received 2 points for a high confidence-correct choice, 1 point for a low confidence-correct choice, 0 points for a low confidence-incorrect choice, and lost 1 point for a high confidence-incorrect choice. The motion animation and the experiment interface were implemented in Python 3.7 (mainly Matplotlib 3.1.0) and displayed at 50 fps with the Qt 5.12 backend. Sessions took ca. 70 minutes per participant. All participants provided informed consent at the beginning of the study. The experiment was approved by the Harvard Institutional Review Board (IRB15-2048). No data was excluded from the analysis.

#### Experiment 1: 4-AFC visual motion structural inference task

The four motion structures used during training and the subsequent experiment were specified by their motion strengths:

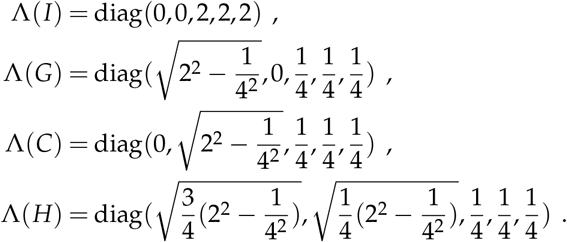

We had chosen these strength values based on pilot experiments with the aim to provoke a significant fraction of error trials while not risking task engagement. During training, participants performed ~30 trials for which the true structure of the upcoming trial was revealed by showing the structure’s name (see **Supplemental Video 1** for structure names) at the center of the screen at the beginning of the trial, and participants could ask for more training trials until they felt ready to start the experiment. During the experiment, participants were asked to identify the motion structure underlying 200 trials (as sketched in **Figure 2A**): per participant, 100 unique trials were generated by uniformly sampling 1 of the 4 motion structures. Each trial was duplicated, and the resulting 200 trials were presented in shuffled order. Participants showed no awareness of trial repetitions during debriefing. The motion structure was drawn randomly before each trial and remained fixed thereafter. After the participant had made a choice, the button turned green if the choice was correct. If incorrect, the button turned red and the correct answer was highlighted in green.

#### Experiment 2: ambiguous 2-AFC visual motion structural inference task

The same participants performed the ambiguous 2-AFC part, following the 4-AFC part after a short break. They were instructed to distinguish between *C* and *H*, because these two structures are particularly hard to distinguish (as sketched in **Figure 3A**). Unbeknown to the participants, we morphed 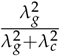 between 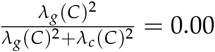 and 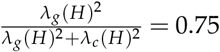, creating ambiguous, intermediate structures between the prototypes *C* and *H*. Each participant received 100 2-AFC trials with 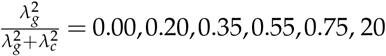 trials each, presented in shuffled order. Due to the limited number of trials, we refrained from trial repetitions in Experiment 2. No feedback was provided at trial end. The explanation given to the participants for removing feedback was that we wanted to avoid any further learning. Due to the lack of a “correct” *C*/*H* answer for most of the trials, the participant’s monetary bonus was calculated based on the choice that an Bayesian ideal observer (see below) would have made.

### Model details

#### Bayesian ideal observer

The ideal observer treats the dynamics of Equation 2 as latent. Only noisy observations, ***x***(*t*), of the true dot locations, ***z***(*t*), can be observed at the times of the video frames. Velocity is unobservable and must be inferred from the observed locations. Observable locations are modelled as centered at the true locations, ***z***(*t*), yet corrupted by independent Gaussian noise: 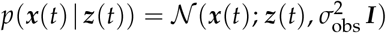, with ***I*** denoting the identity matrix. By inverting this dynamical generative model, we identify Kalman filtering (Bishop, 2006; Kalman, 1960) as the correct method for on-line inference. Using the standard terminology for Kalman filtering, the state transition matrix ***F*** and the covariance of the process noise ***Q*** are given by:

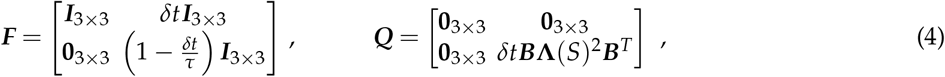

with *δt* denoting the time between consecutive frames. On a cautionary note, Kalman filters employ a Gaussian posterior distribution which, from a strict standpoint, is not compatible with the circle. However, all dots are always visible and, thus, the uncertainty about location in the posterior is much smaller than *π*. Consequently, inference errors due to circularity of the stimulus are negligible.

Kalman filters provide an efficient method for calculating the likelihood of a presented trajectory recursively from the pre-fit residuals, via 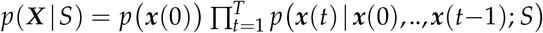. In the numerical evaluation of trajectory likelihoods, we set ***x***(*t*) to the dot locations of the presented trial, without applying noise to the presented locations. This choice was made for computational reasons because, when applying noise, we would have to evaluate many trial repetitions (with different noise realizations) to obtain a stable estimate of the trajectory likelihood. Nonetheless, we decided to model observation noise in the observer (the inference machine) for biological realism. Without observation noise, the Kalman filter could base its decision on minuscule correlated velocity changes at single-frame precision—an unlikely faculty for any biological observer. The value of *σ*_obs_ was determined by maximum likelihood fitting, as explained below in “Model fitting”.

A technical complication, that we have not discussed so far, lies in the assignment between dot identities (their role in the structure) and dot colors (colors were randomized in the stimulus). Using the example of clustered (*C*) motion, grouping of any two of the three dots is a valid form of the clustered motion structure and leads to the same response (choice ‘*C*’). However, the Kalman filter calculates the trajectory likelihood for only one particular assignment. For clarity of the presentation, we had left out this detail in the Results section. Here, we present the full ideal observer which accounts for the multiplicity of dot assignments. We accommodate dot assignments by means of three dot permutations,

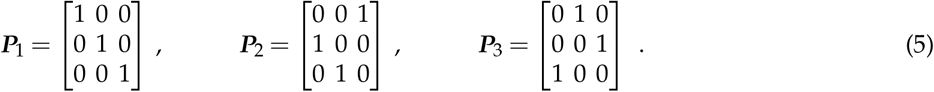

Returning to the example of *C* choices, we need three separate Kalman filter evaluations, calculating *p*(***X*** | ***P**_m_****BΛ***(*C*) for *m* = 1,2,3, in order to accommodate all possible clustered structures. We denote the set of involved permutations by multiplicity *M*(*C*) = {1,2,3}. Similarly, hierarchical structure also has a multiplicity of |*M*(*H*)| = 3 with *M*(*H*) = {1,2,3}. In contrast, independent motion and global motion are permutation-invariant, and we have *M*(*I*) = *M*(*G*) = {1}. Incorporating the structure multiplicities, we obtain:

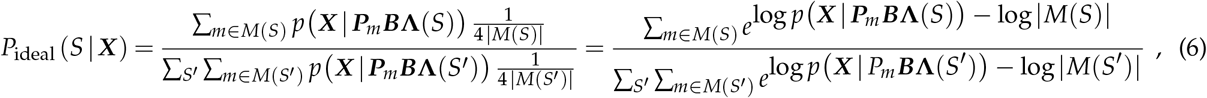

where log denotes the natural logarithm, and the multiplicity penalty |*M*(*S*)| ensures that the posterior distribution in the absence of evidence matches the optimal prior, 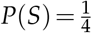 for all *S*. The last equality is not required for evaluating the ideal observer’s posterior, but will ease the understanding of the choice model discussed next.

#### Choice model

The choice model augments the ideal observer with three known human imperfections. First, human participants may be biased towards certain structures. This is accounted for by a prior probability *P*(*S*) for perceiving each of the four structures. Since the prior must sum to one, this leaves three degrees of freedom such that we can express the log-prior 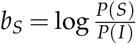 relative to independent motion, for *S* = *G, C, H*. Second, human choices often are not deterministic, but stochastic. We model choice stochasticity by exponentiating the posterior with inverse temperature parameter, 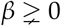. For *β* → ∞, we recover deterministic selection of the option with the highest posterior belief; for *β* → 0, we obtain uniformly random choices. Third, in some trials, human choices cannot be explained by a simple rational model (for example, the participant may have been inattentive during stimulus presentation). To account for these trials, we adopt the common method of including a lapse probability, *π_L_*, that a uniformly random response is given. The resulting model is a “stochastic posterior decision maker with lapses and imperfect prior”, the winning model in the systematic model comparison by Acerbi, Vijayakumar, and Wolpert, (2014), and has six parameters (*σ*_obs_, *τ_L_, β, b_G_, b_C_, b_H_*).

As for the ideal observer, we had not discussed the technical complication of structure multiplicities, *M*(*S*), in Equation 1 of the Results section. Including the structure multiplicities, we obtain:

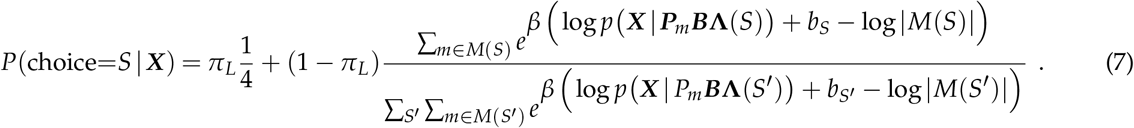

Equation 7 is the equation that we used for model fitting and choice predictions. Likewise, when evaluating the Bayesian decision confidence, the x-axes of **Figure 2I** and **Figure 3D** respect structure multiplicities by setting *π_L_* = 0 and *β* = 1 in Equation 7.

### Model fitting and prediction of human choices

#### Model fitting via maximum likelihood

The psychometric curve of the choice model can be horizontally stretched by *σ*_obs_, which injects noise into evidence accumulation, or vertically squeezed by *π_L_*, which narrows the range of the choice distribution. Thus, both *σ*_obs_ and *π_L_* overlap with *β* in controlling the slope of the curve. To make the model more identifiable, we fitted *σ*_obs_ and *π_L_* as shared values for all participants. Denoting by *S*^(*j,i*)^ the choice participant *j* made in the *i*-th trial ***X***^(*j,i*)^, the maximum likelihood values of the shared parameters were obtained via grid search for computational efficiency:

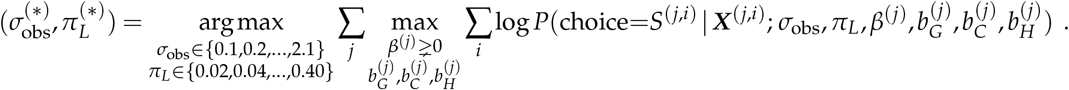

We find 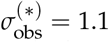 and 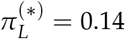 for the 4-AFC task. We maintained these values also for the 2-AFC task because of the larger data corpus of Experiment 1, and for better comparability of parameters across experiments. Furthermore, refitting *σ*_obs_ and *π_L_* to the 2-AFC data did not significantly improve the likelihood of human choices under the model.

Then, we fitted the remaining free parameters to each participant *j* via maximum likelihood:

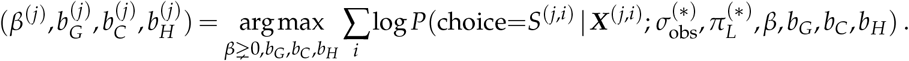

For the 2-AFC experiment, since *I* and *G* are not present in the task, we removed all *I*- and *G*-related terms from Equation 7 and set *b_C_* = 0 as the reference bias, leaving *β* and *b_H_* as the only two free parameters to fit.

#### Cross-validated prediction of choices

Unless noted otherwise (see next paragraph), we used leave-one-out cross-validation to generate model predictions: the predicted choice distribution, *p*(choice = *S* | ***X***^(*j,i*)^), to trial *i* was obtained by fitting the per-participant parameters to all other trials of the *j*-th participant in the respective experiment. Also, all log-likelihood comparisons are based on the cross-validated predicted choice distributions.

There were two exceptions to this evaluation. First, for the cross-participant model evaluation in **Supplemental Figure S4**, we fitted the per-participant parameters to the entire 4-AFC dataset of each participant, and evaluated each fitted model on the entire 4-AFC dataset of every participant. Second, for the transfer model in **Figure 3**, we fitted 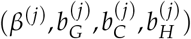 to the entire 4-AFC dataset of participant *j*, and used 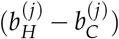 as the bias for the 2-AFC predictions, as the relative log-prior between *H* and *C* can be decomposed as 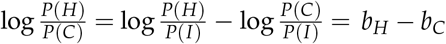.

### Details on the non-Bayesian choice model

The non-Bayesian model compares the dot velocity correlation in the presented trial to the prototypes of the different structures, thereby converting structural inference into a clustering task. Concretely, let ***ρ***^(*j,i*)^ denote the three pairwise Pearson correlation coefficients,

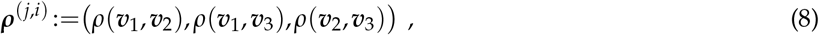

between the noise-free, empirical velocities for the *i*-th trial of the *j*-th participant, obtained from subsequent stimulus frames, 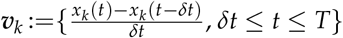. While stripped of dynamic transitions between frames, velocity correlations are still an informative cue about the underlying motion structure. For example, global (*G*) motion is characterized by strong positive correlations between all dots’ velocities. For the prototypes of all motion structures, the correlations ***ρ***(***P***_*m*_**BΛ**(*S*)) between dot velocities are given by the off-diagonal elements of 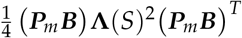, according to Equation 3. Hereby, we respect the multiplicities, *M*(*S*), of the different structures. Examples of the correlations from presented trials as well as the prototypes are shown in **Supplemental Figure S8**.

The resulting non-Bayesian model retains the architecture of Equation 7, but replaces the ideal observer’s log-likelihood by the correlational distance:

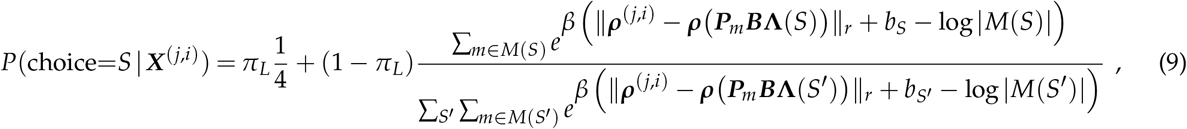

with ║·║_*r*_ denoting the *r*-norm. Like the full choice model with Bayesian core, it has 6 parameters (*r* replaces *σ*_obs_). For fair comparison between the Bayesian and non-Bayesian models, we fitted *π_L_* and *r* as shared parameters across participants, and fitted the remaining 4 parameters to each individual. For the 4-AFC task, we obtained *π_L_* = 0.30 and *r* = 1.0.

## Code and data availability

The Python code for data collection and analysis, as well as the de-identified data, is available online at https://github.com/DrugowitschLab/motion-structure-identification.

## Acknowledgments

We thank Eric Schulz for helpful feedback and discussions at multiple stages of the research. The work was supported by grants from the NIH (R01MH115554, J.D.), the Harvard Brain Science Initiative (Collaborative Seed Grant, J.D. & S.J.G.), the Center for Brains, Minds, and Machines (CBMM; funded by NSF STC award CCF-1231216), an Alice and Joseph Brooks Fund Fellowship award (J.B.), and a James S. McDonnell Foundation Scholar Award for Understanding Human Cognition (grant# 220020462, J.D.).

## Supplemental figures

**Supplemental Figure S1.**
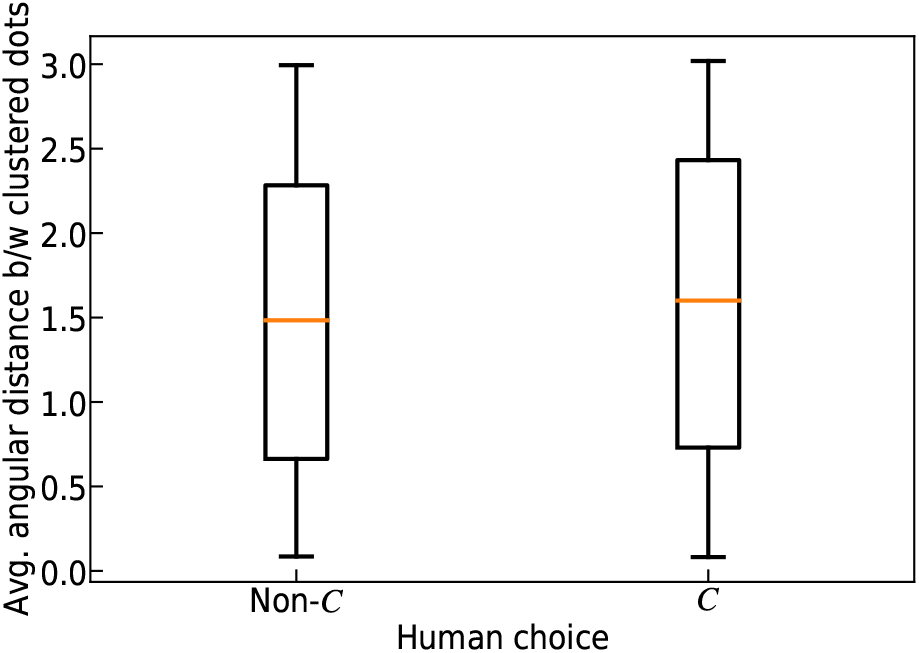
Spatial arrangement of dots could influence motion structural inference. We evaluated, for all trials with clustered (C) motion, the average angular distance between the two clustered dots, conditioned on whether participants reported clustered motion or not. The similarity of dot proximity distributions indicates that participants did not rely on dot proximity when solving the task.

**Supplemental Figure S2.**
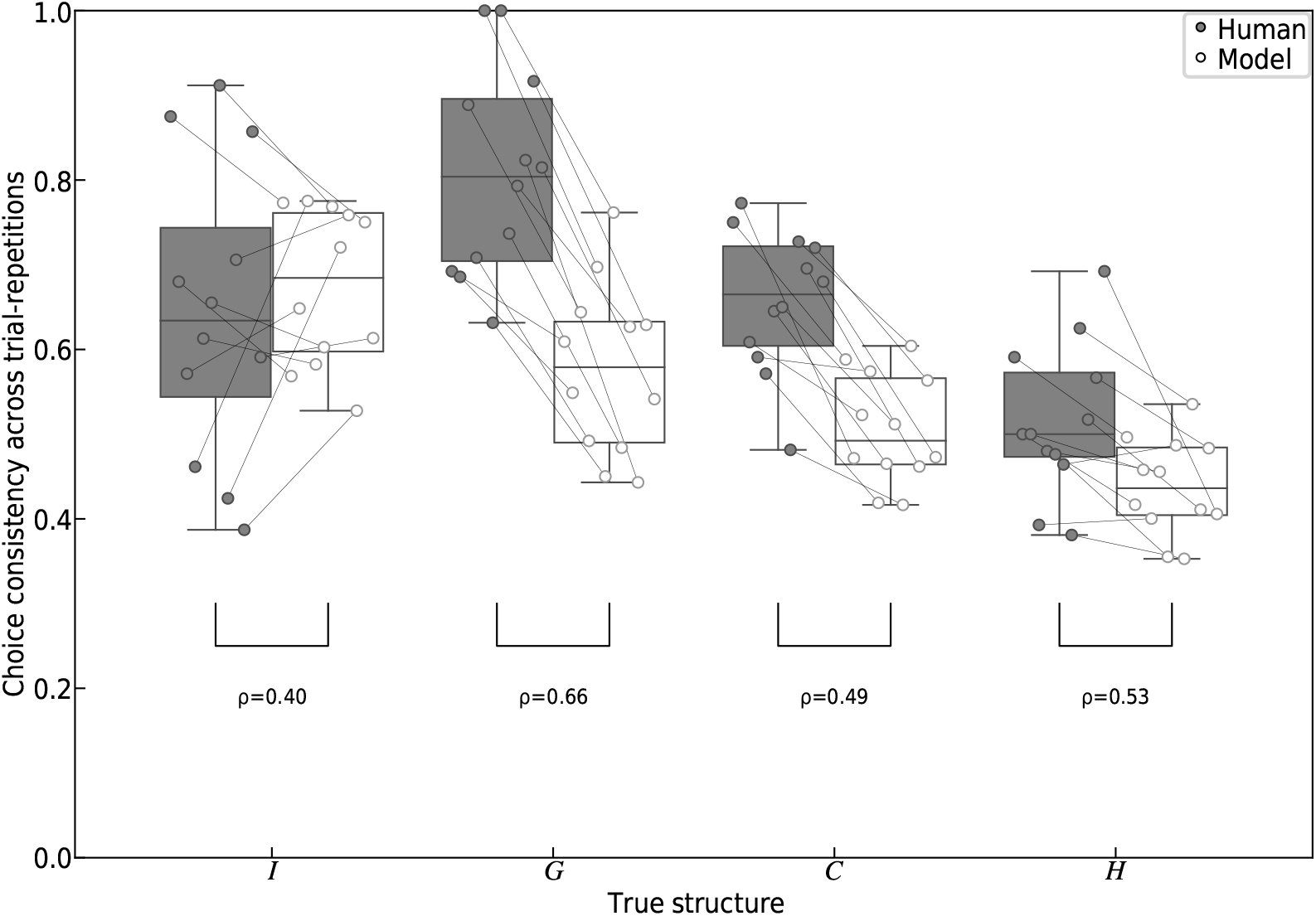
Average consistency of human choices and respective model predictions across repetitions of trials, for each of the four structures. Within each structure, the lines link the human and model consistency of each participant and reveal their correlation. This is quantified by the Spearman’s rank correlation coefficients *ρ* in the figure. The model had a tendency to underestimate human consistency, but qualitatively predicted the pattern across conditions. Box plots mark the 0, 25, 50, 75, and 100 percentiles.

**Supplemental Figure S3.**
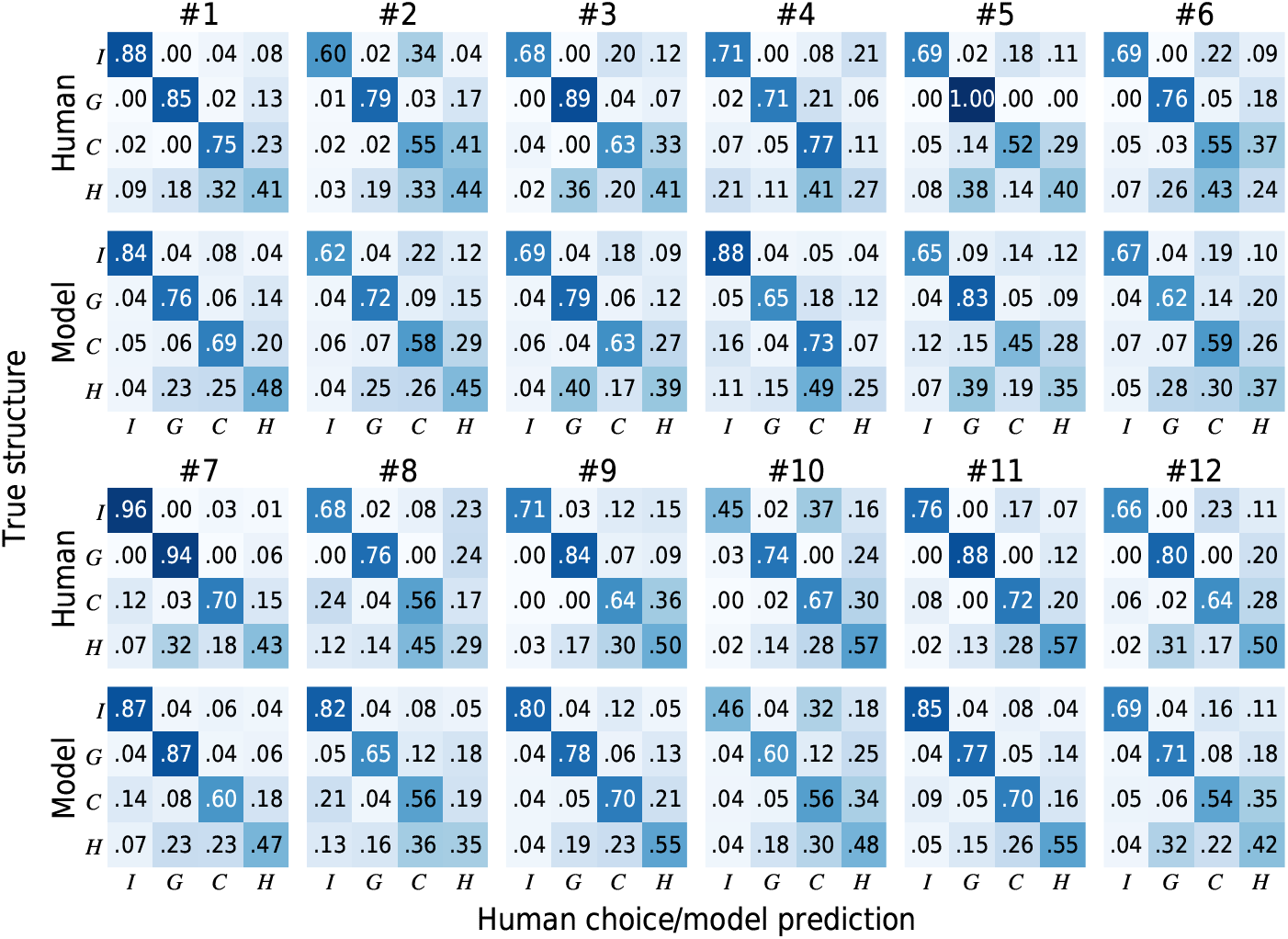
Confusion matrices of all twelve participants and their respective model predictions. Participants featured diverse error patters. For example, some participants confused H-G more than H-C, while others showed the opposite. The choice model captured the full variety of error patterns.

**Supplemental Figure S4.**
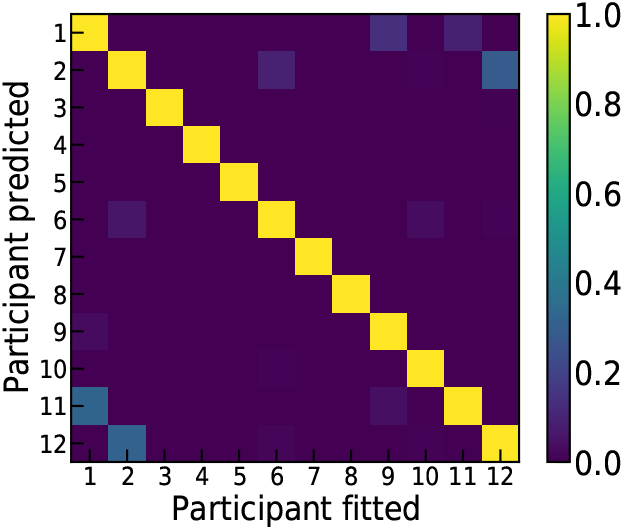
Cross-participant model comparison. The (*i,j*) element is the likelihood of responses of participant *i* under the parameters fitted to participant *j*, normalized to 1 within each row. The dominance of diagonal elements implies that individual differences in the confusion matrices were not driven by the set of presented trials.

**Supplemental Figure S5.**
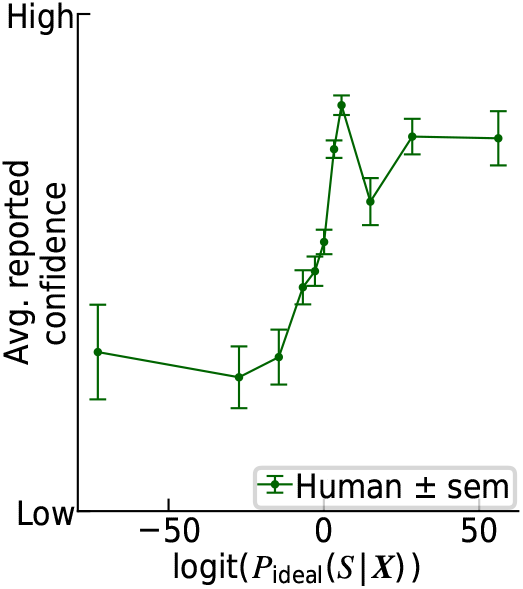
Fraction of human high confidence reports as a function of the ideal observer’s log-odds, logit (*P*_ideal_ (*S*|**X**)). Without including the fitted prior (biases *b_S_*), the curve still exhibits an approximately monotonic increase, similar to **Figure 2I**.

**Supplemental Figure S6.**
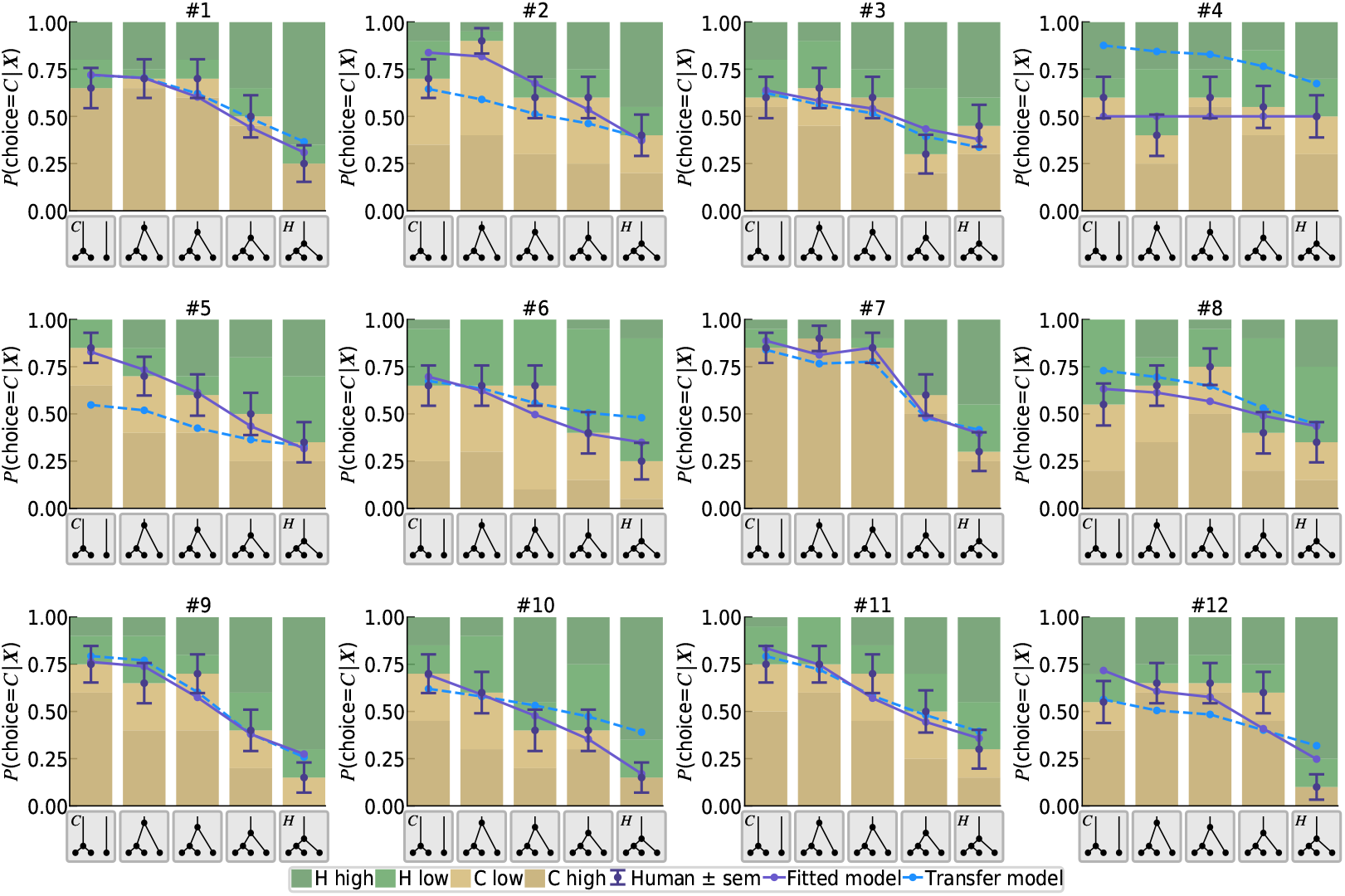
Same as **Figure 3B**, but for every participant separately. The fitted model (solid purple line) captured the diverse human transitions of choice, as the structure morphed from *C* to *H*. Also, the transfer model (dashed blue line) predicted many participants’ responses accurately.

**Supplemental Figure S7.**
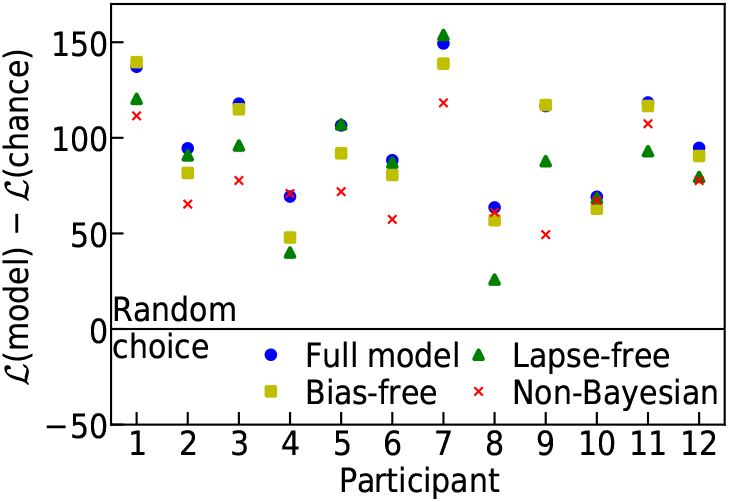
Model comparison in terms of the log-likelihood of human responses under the different choice models of **Figure 4A**. All log-likelihoods are evaluated relative to chance level (uniform choice with probability 1/4).

**Supplemental Figure S8.**
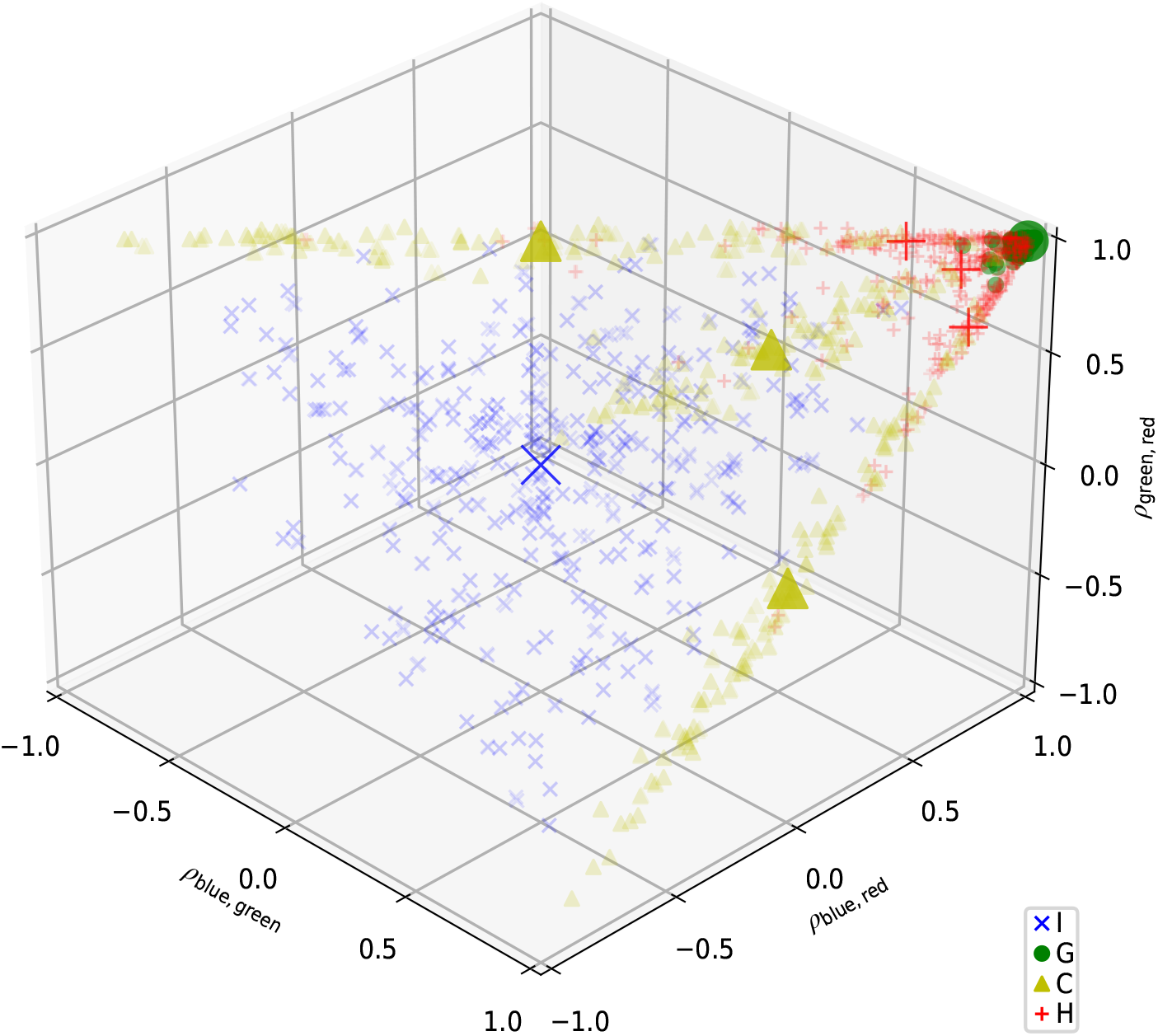
Distribution of the three Pearson correlation coefficients (*ρ*_blue,green_,*ρ*_blue,red_,*ρ*_green,red_) of the empirical dot velocity trajectories ***V***(***X***). Colors and markers indicate the generating trial structure. All 1200 unique trials of Experiment 1 are shown. The larger symbols mark the prototype of the motion structures (including multiplicity for *C* and *H*). See Materials and Methods for mathematical details about the prototypes and their multiplicity. Trials of the same structure tend to cluster around their prototype, such that decision boundaries could be drawn between different structures, laying the basis for the non-Bayesian, prototype-based model of motion structural inference.

